# Biomolecular Condensates defined by Receptor Independent Activator of G protein Signaling: Properties and Regulation

**DOI:** 10.1101/2023.05.02.539089

**Authors:** Ali Vural, Stephen M. Lanier

**Author notes:** Corresponding author Ali Vural, Ph.D. Present address: Department of Biomedical Sciences Western Michigan University Homer Stryker M.D. School of Medicine 1000 Oakland Dr, Kalamazoo, MI 49008.

## Abstract

Activator of G-protein Signaling 3 (AGS3), a receptor independent activator of G-protein signaling, oscillates among different subcellular compartments in a regulated manner including punctate entities referred to as biomolecular condensates (BMCs). The dynamics of the AGS3 oscillation and the specific subcompartment within the cell is intimately related to the functional diversity of the protein. To further address the properties and regulation of AGS3 BMCs, we asked initial questions regarding a) the distribution of AGS3 across the broader BMC landscape with and without cellular stress, and b) the core material properties of these punctate structures. Cellular stress (oxidative, pHi, thermal) induced the formation of AGS3 BMCs in two cell lines (Hela, COS7) as determined by fluorescent microscopy. The AGS3-BMCs generated in response to oxidative stress were distinct from stress granules (SG) as defined by the SG marker protein G3BP1 and RNA processing BMCs defined by the P-body protein Dcp1a. Immunoblots of fractionated cell lysates indicated that cellular stress shifted AGS3 to the membrane pellet fraction, whereas the protein markers for stress granules (G3BP1) SG- BMCs remained in the supernatant. We next asked if the formation of the stress-induced AGS3 BMCs was regulated by protein binding partners involved with signal processing. The stress-induced generation of AGS3 BMCs was regulated by the signaling protein Gαi3, but not by the AGS3 binding partner DVL2. Finally, we addressed the fluidity or rigidity of the stress-induced AGS3-BMCs using fluorescent recovery following photobleaching of individual AGS3-BMCs. The AGS3-BMCs indicated distinct diffusion kinetics that were consistent with restricted mobility of AGS3 within the stress-induced AGS3-BMCs. These data suggest that AGS3 BMCs represents a distinct class of stress granules that define a new type of BMC that may serve as previously unappreciated signal processing nodes.

**Summary statement:** AGS3 assembles into distinct biomolecular condensates in response to cell stress and this assembly is selectively regulated by AGS3 binding partners involved in signal transduction within the cell.

## INTRODUCTION

Activators of G protein Signaling (AGS) proteins define a broad panel of biologic regulators that influence G protein signaling, however, how this regulation mechanistically integrates signals within different systems for physiological system homeostasis and in the context of pathophysiological dysfunction is not fully understood. One member of the AGS family of particular interest is AGS3 (G-protein signaling modulator 1, *Gpsm1*), a Group II AGS protein. AGS3 is implicated in a broad range of functional roles in specific cell types, tissues and organ systems (Vural et al., 2010, Oner et al., 2013b, Vural et al., 2013, Vural and Lanier, 2020, Vural et al., 2019, Vural et al., 2018, Vural et al., 2016, Huang et al., 2014, Oner et al., 2010, Nadella et al., 2010, Regner et al., 2011, Kwon et al., 2012, Adekoya et al., 2019, Blumer et al., 2008, Rasmussen et al., 2015, Yip et al., 2020, Sato et al., 2004, Xing et al., 2015, Nie et al., 2020, Pattingre et al., 2004, Williams et al., 2011, Gotta et al., 2003, Kamakura et al., 2013, Branham-O’Connor et al., 2014, Zhao et al., 2010, Blumer et al., 2002, Sanada and Tsai, 2005, Pattingre et al., 2003, Garcia-Marcos et al., 2011, Xu et al., 2010, Yeh et al., 2013, Wang et al., 2013, Groves et al., 2010, Blumer et al., 2003, An et al., 2008, Oner et al., 2013a, Groves et al., 2007), however it has been challenging to define the mechanistic elements of the various functional roles as mentioned above. AGS3 exhibits a tissue-specific expression pattern that is regulated during development and during tissue response or adaptation to various physiological or pathophysiological challenges. (Bowers et al., 2004, Yao et al., 2005, Bowers et al., 2008, Blumer et al., 2008, Fan et al., 2009, Nadella et al., 2010, Regner et al., 2011, Kwon et al., 2012, Wang et al., 2013). AGS3 actually oscillates between cytosolic and membrane distribution moving among different subcellular compartments (e.g. cytosol (Pizzinat et al., 2001), cell membrane (Oner et al., 2010, Oner et al., 2013a, Robichaux et al., 2015), biomolecular condensates (Vural et al., 2010, Vural et al., 2018, Vural and Lanier, 2020), centrosome (Blumer et al., 2006), *trans*-Golgi network of the Golgi Apparatus (Oner et al., 2013b, Nie et al., 2020), and the aggresomal pathway (Vural et al., 2010) in a regulated manner. One premise of our recent work in this area is that the understanding of the factors regulating the movement of AGS3 among these different subcellular compartments will provide critical insight to the functional mechanisms mentioned above.

AGS3 has seven N-terminal tetratricopeptide repeats (TPR) and four C-terminal G-protein regulatory (GPR) motifs connected with a linker domain. TPR repeats influence intramolecular dynamics and subcellular distribution of AGS3 through interaction with specific binding partners. The GPR motifs in AGS3 act as guanine nucleotide dissociation inhibitors (GDI) stabilizing the Gα subunit in its GDP-bound conformation free of Gβγ (Peterson et al., 2000, Pizzinat et al., 2001, Bernard et al., 2001) and this interaction also influences subcellular distribution of AGS3.

Of particular interest, AGS3 appears to have an inherent propensity to form distinct, non-membranous punctate structures (<200 nm to 2.5 μm) in multiple cell types (Vural et al., 2010, Vural et al., 2018) Single amino acid substitutions within AGS3 lead to dramatic appearance of these punctate entities, which are apparently not associated with defined vesicle or organelle markers (Vural et al., 2010, Vural et al., 2018) and are henceforth referred to as AGS3 biomolecular condensates (BMCs) (Vural and Lanier, 2020). BMCs are defined as non-membranous and micron-scale compartments in eukaryotic cells that are formed upon phase separations (i.e., liquid-liquid phase separation, LLPS, or phase transitions) (Banani et al., 2017, Glauninger et al., 2022). BMCs provide a platform to coordinate the assembly of a specific subset of molecules sequestered from the rest of the cytoplasm to facilitate a wide range of biological and chemical events within the cell (Mitrea et al., 2022). Key parameters driving BMC formation include intrinsically disordered regions (IDRs) within any given protein and the apparent multivalent character of its interaction with various binding partners, both of which are characteristic properties of AGS3 (GPSM1) and the related protein AGS5 (LGN, GPSM2) (Sato et al., 2006, Blumer and Lanier, 2014, Uversky, 2014, Erdos and Dosztanyi, 2020). The formation of any given BMC may be regulated by cell stimuli and various BMCs play various roles in cell signal amplification, storage and trafficking of biomolecules, and/or the concentration of biochemical reactions (Mitrea et al., 2022).

We previously identified the regulated engagement of AGS3 with Dishevelled-2 (DVL2) signalosomes or DVL2 BMCs. In various contexts, DVL2 functions as a “hub” or “anchor” protein coordinating Wnt signaling integration (Turm et al., 2010, Sharma et al., 2018, Vural and Lanier, 2020). Of note, DVL2 and AGS3 are involved to varying degrees with similar tissue and cell functions and adaptations including cilia formation (Yeh et al., 2013, Lee et al., 2012, Xie et al., 2018), cell polarity (Blumer et al., 2003, Gotta et al., 2003, Izaki et al., 2006, Saadaoui et al., 2017, Sharma et al., 2018, Zhang et al., 2007) and system adaptations associated with addiction (Bowers, 2010, Dias et al., 2015). AGS3 is apparently “recruited” to DVL2 puncta (Vural et al., 2018) and interacts with DVL2 in a phosphorylation dependent manner. Moreover, the distribution of AGS3 to DVL2 BMCs is regulated by a cell-surface, G-protein coupled receptor (Vural et al., 2018, Vural and Lanier, 2020). Interestingly, however, phosphorylation-deficient AGS3 (AGS3-T602A) and AGS3 TPR point mutants exhibited punctate structures that seem distinct from DVL2 signaling puncta (Vural et al., 2018).

This manuscript is part of an expanded focus on the biology and regulation of AGS3 BMCs and the concept of “image phenotype profile screens” or “image-signaling system connectivity” as mentioned in an earlier report (Vural and Lanier, 2020). In the current study we addressed a) the distribution of AGS3 to defined BMCs, b) the influence of cell stress on BMC generation and its interaction with various binding partners and c) the mobility or diffusion kinetics for AGS3 in the context of distinct BMCs. The results of these studies suggest that AGS3 BMCs, the formation of which is regulated by both cellular stress and the signaling proteins Gαi3 and DVL2, define a new type of BMC that may serve as previously unrecognized signal processing nodes.

## MATERIALS AND METHODS

### Materials

Sodium (meta)arsenite (NaAsO2, S7400), beta-Actin-Peroxidase antibody (mouse monoclonal, A3854), diamide (D3648) and 2-Mercaptoethanol (M7154) were purchased from MilliporeSigma (Burlington, MA). AGS3 antibody (mouse monoclonal [G-2], sc-271721), DVL2 antibody (mouse monoclonal [10B5], sc-8026), eukaryotic initiation factor 4 gamma (eIF4G) antibody (mouse monoclonal [A-10], sc-133155) and mRNA-decapping enzyme 1A (Dcp1a) antibody (mouse monoclonal [56-Y], sc-100706) were purchased from Santa Cruz Biotechnology (Santa Cruz, CA). Polyethyenimine (PEI) (linear, *M*r∼25,000) was obtained from Polysciences, Inc. (Warrington, PA). DC Protein Assay Kit II (5000112) was purchased from BioRad (Hercules, CA). Halt^TM^ Protease and Phosphatase Inhibitor (1861280), Blocker^TM^ BSA in TBS (37520), Spectra^TM^ Multicolor Broad Range Protein Ladder (26634), SuperSignal West Dura Extended Duration Substrate (34075), Prolong Diamond Antifade (P36961) reagent, Novex WedgeWell 4-20% Tris-Glycine Gels (XP04202BOX), iBlot Gel Transfer Stacks PVDF - regular (IB401001) and iBlot Gel Transfer Device (IB1001), DMEM (21063- 029), Hank’s Balanced Salt Solution (HBSS; 14025-076 including calcium chloride (CaCl2) (anhydrous.), magnesium chloride (MgCl2-6H2O), magnesium sulfate (MgSO4-7H2O), potassium chloride (KCl), potassium phosphate monobasic (KH2PO4), sodium bicarbonate (NaHCO3), sodium chloride (NaCl), sodium phosphate dibasic (Na2HPO4) anhydrous and D-Glucose (Dextrose)) and HBSS (14185-052 including potassium chloride (KCl), potassium phosphate monobasic (KH2PO4), sodium chloride (NaCl), sodium phosphate dibasic (Na2HPO4-7H2O), D-Glucose (Dextrose)) were purchased from Thermo Fisher Scientific (Waltham, MA). pRC/CMV::DVL-2-Myc was obtained from Addgene (Cambridge, Massachusetts, USA) (plasmid #42194) as deposited by Robert Lefkowitz and Shin-ichi Yanagawa (Lee et al., 1999). pT7-EGFP-C1-HsDCP1a was a gift from Elisa Izaurralde (Addgene plasmid # 25030; http://n2t.net/addgene:25030;RRID:Addgene_25030) (Tritschler et al., 2009). Ras GTPase-activating protein-binding protein 1 (G3BP1) phage UbiC G3BP1-GFP-GFP was a gift from Jeffrey Chao (Addgene plasmid # 119950; http://n2t.net/addgene:119950;RRID:Addgene_119950) (Wilbertz et al., 2019). All other materials were obtained as previously described.

### Cell Culture, Cellular Transfection, Immunoblotting and Fractionation

COS-7 (CRL-1651) and HeLa (CCL-2) cell lines were purchased from American Type Culture Collection (ATCC) (Manassas, VA). Both cell lines were cultured in Dulbecco’s Modified Eagle Medium (high glucose) supplemented with 2 mM glutamine, 100 units/ml penicillin, 100 μg/ml streptomycin and 10% fetal bovine serum. COS-7 cells were transfected using PEI as described previously (Vural et al., 2018). To generate AGS3-expressing stable HeLa cell lines, wild-type AGS3-GFP plasmid was transfected into HeLa cells, using Lipofectamine 2000 as previously described (Vural et al., 2016). The cells were selected with 1000 μg/ml G418 and sorted based on GFP expression.

For immunoblotting and fractionation, cells were lysed with a buffer (250 μl) consisting of 25 mM HEPES (pH 7.4), 4% glycerol, 0.5% (v/v) Nonidet P-40, 150 mM NaCl, 2 mM CaCl2 and Halt^TM^ Protease and Phosphatase Inhibitor (Thermo Scientific, Waltham, MA). Lysates were shaken on ice for 15-20 minutes followed by centrifugation at 13,000 rpm for 10 minutes at 4°C. The supernatant was transferred and protein concentration was determined using DC Protein Assay Kit II (BioRad, Hercules, CA). The pellet was resuspended in 100 μl of lysis buffer including DNase (10 U/ml) and incubated 30 minutes at room temperature. Immediately after cell lysis and fractionation, the samples (e.g., 25% of the supernatant fraction and 50% of the pellet fraction) were processed by denaturing polyacrylamide gel electrophoresis (SDS-PAGE) using Novex 4-20% gradient gels (Thermo Fisher Scientific, Waltham, MA, USA). For SDS-PAGE with standard reducing agents to disrupt sulfydryl bonds, protein loading buffer (25 mM Tris, pH=6.8; 5% glycerol; 1% SDS; 0.2% bromophenol blue) included 2-Mercaptoethanol. For SDS-PAGE under non-reducing conditions, protein loading buffer (25 mM Tris, pH=6.8; 5% glycerol; 1% SDS; 0.2% bromophenol blue) lacked 2-Mercaptoethanol (2.5%). Protein loading buffer (10 μl) was added to the samples (90 μl) and heated for 10 min at 70 °C. The proteins were transferred onto the PVDF membrane with P3 program for 10 minutes (20V) using the iBlot Gel Transfer Stacks. The membranes were then processed for immunoblotting as previously described (Vural et al., 2018).

### Fluorescence confocal microscopy, Fluorescence Recovery After Photobleaching (FRAP) assays and image analysis

HeLa and COS-7 cells were processed for immunofluorescent microscopy as described (Vural et al., 2018, Vural and Lanier, 2020) and cell images captured with a 63× oil immersion objective on a Zeiss LSM 800 confocal microscope (Microscopy, Imaging & Cytometry Resources Core at Wayne State University). For FRAP assays, glass-bottom dishes with COS-7 cells were placed into the environmental chamber (37 °C, 5% CO2) of the Zeiss LSM 800 confocal microscope. Using bleaching and time series options in ZEN software, individually selected AGS3 puncta were bleached (i.e., fluorescence intensity was nullified) and the images were captured at every three seconds with a total of 40 cycles. The fluorescence intensity of the AGS3 puncta was measured in each image, which was captured at every three seconds. The fluorescence intensity was normalized by subtracting the post-bleaching intensity (t=12 sec) from the pre-bleaching intensity (t=0 sec) for AGS3 puncta from different experiments. All images were obtained from approximately the middle plane of the cells and images were visualized and evaluated through the Adobe Photoshop CC 2018 platform.

### Statistical analysis

Biomolecular condensates generated by cell stress exhibit particle sizes distributing between 0.1-0.3 μm in individual cells. For each experiment, BMCs were counted in at least 200 individual cells to determine the % of cells containing stress granules and AGS3 puncta. Data are expressed as mean ± SEM as determined from at five or more independent experiments. For photobleaching experiments, data are expressed as mean ± SEM as determined from five or more independent experiments. Five to ten individual AGS3 puncta from multiple individual COS-7 cells were photobleached to determine the fluorescence intensity in each individual experimental group. Data at time t=120 s after bleaching were analyzed with Prism for Mac OS X (Version 9.1.2) software (GraphPad Software, San Diego, CA) using one-way *ANOVA*, where significant differences between groups were determined by Tukey’s multiple comparison test. *P-*values were calculated on t=120 s comparable time point and *P* values <0.05 were considered statistically significant.

## RESULTS

AGS3 typically exhibits a non-homogeneous cytosolic and membrane distribution in cells, but also assembles into distinct, non-membranous punctate structures and oscillates between various functional microdomains within the cell in a regulated manner (Vural et al., 2010, Oner et al., 2013b, Huang et al., 2014, Vural et al., 2018, Vural and Lanier, 2020). As previously reported, with exogenous expression of AGS3 in various cell types, approximately ±80% of AGS3 expressing cells exhibit a non-homogeneous cytosolic and membrane distribution in cells, whereas ±20% exist as punctate entities. Targeted disruption of the TPR domain of AGS3 and/or altered phosphorylation status of AGS3 appear to direct the protein to such punctate structures, which are not associated with any defined vesicle or organelle markers (Vural et al., 2010) thus exhibiting characteristics of biomolecular condensates.

As a protein, AGS3 exhibits significant regions of intrinsic disorder, which is a property also observed with other proteins that engage with and/or generate biomolecular condensates (Shi et al., 2021). AGS3 possesses two defined multivalent motifs [(7 tetratricopeptide repeats (TPR), 4 G-protein regulatory (GPR) motifs] connected through a linker domain region. We quantified the predicted enrichment of intrinsically disordered regions (IDRs) in AGS3 protein sequence using the IUPred algorithm (Dosztanyi, 2018). Approximately 40% of the AGS3 protein, including the linker and GPR domains, is predicted to exist as IDRs (Figure 1A). It is likely that the GPR domain becomes structurally organized upon its interaction with Gα binding. IDRs typically fluctuate in form and may engage in number of multivalent interactions with varying degrees of affinity as the region oscillates across a spectrum of fluid energy states and surface presentations with and without various protein or lipid binding partners. Disruption of the organizational structure of either the TPR or GPR domains by targeted point mutations also leads to the formation of AGS3 BMCs (Vural et al., 2010, Vural et al., 2018). As observed with oxidative cell stress in the present manuscript, targeted point mutations in the TPR or GPR domains also increase the relative distribution of AGS3 to the membranous pellet following cell lysis (Vural et al., 2010, Vural et al., 2018). Another key feature of the generation of BMCs is within the context of how cells respond to external stressors. We thus determined the impact of three environmental stressors (oxidative, pHi, and thermal) on AGS3-BMCs formation. Strikingly, each environmental stressor resulted in the generation of AGS3-BMCs (Figure 1B). Similar observations were observed with both transfected and endogenous AGS3 (Vural, A and Lanier, SM – unpublished observations).

**Figure 1.**
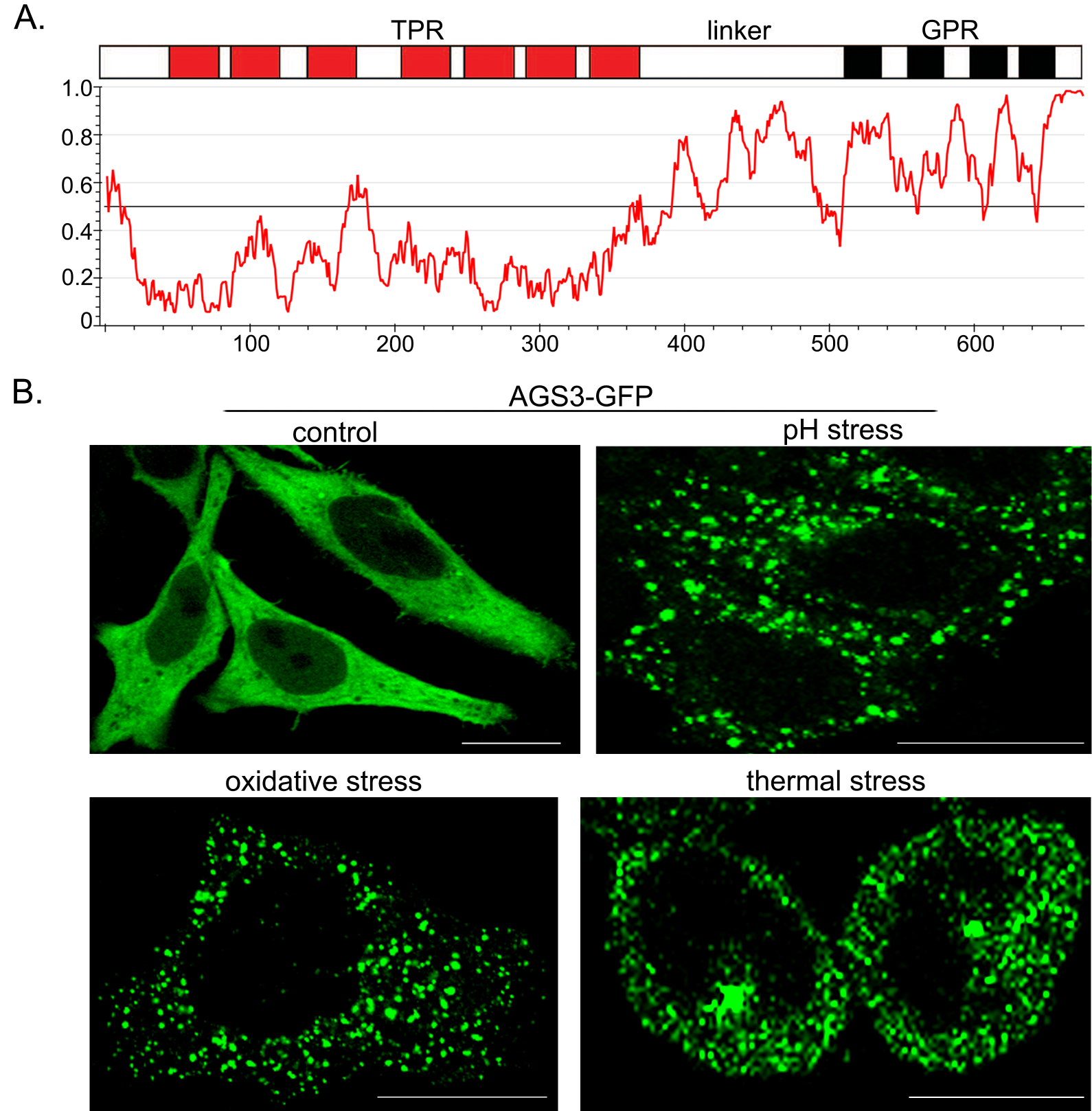
AGS3 domain organization and the influence of environmental stressors on the subcellular distribution of AGS3. A) The disorder score of full length AGS3 amino acid sequence (NCBI genbank # NP_653346, *Rattus norvegicus*) was revealed by IUPRED2 (Dosztanyi, 2018), a prediction server of intrinsically unstructured regions of proteins based on estimated energy content. Probability values over 0.5 (y-axis) are considered as disordered. B) HeLa cells stably expressing AGS3-GFP were exposed to HBSS without bicarbonate (pH stress; 30 min), thermal stress (T=42.5°C; 1 hour), and oxidative stress (arsenate; 500 nM; 30 min) and processed for immunocytochemistry. Scale bars: 10 μm.

In our previous studies, we also reported the engagement of wild type AGS3 with DVL2 signaling BMCs which is regulated by DVL2 phosphorylation, G-proteins and cell surface G-protein coupled receptors (Vural et al., 2018, Vural and Lanier, 2020). We thus extended this work to investigate the engagement of AGS3 with other defined BMCs. As an initial approach, we determined the cellular distribution of AGS3 and BMC biomarkers for the RNA processing BMCs generated in response to oxidative cellular stress (sodium arsenate) (Mittal and Flora, 2006, Jiang et al., 2013) in two distinct cell types – a human epithelium cell line (HeLa) and the primate renal fibroblast-like cell line (COS-7). Stress granule BMCs are non-membranenous compartments consisting of RNA and RNA-binding proteins (RBPs) that are formed as a response to certain stressors such as oxidative cell stress or heat shock (Youn et al., 2019, Glauninger et al., 2022). Stress granules are one of the first defined BMCs that appear impact mRNA translation and stability and have been linked to the pathogenesis of a closely related set of degenerative diseases, including amyotrophic lateral sclerosis, frontotemporal dementia, and inclusion body myopathy (Molliex et al., 2015, Wolozin and Ivanov, 2019). Stress granules are typically identified with both an endogenous biomarker (eIF4G) and an exogenously expressed biomarker (G3BP1) and their formation is induced in response to various types of cell or tissue stress (Kedersha and Anderson, 2007, Molliex et al., 2015).

### Cellular distribution of AGS3, stress granule proteins and P-body proteins with and without oxidative cell stress

In non-stressed cells, AGS3 and the SG BMC marker proteins eIF4G and G3BP1 exhibited a non-homogenous cytosolic or membrane distribution in both Hela and COS7 cells (Figure 2A-B). The cellular distribution of the SG BMC marker proteins eIF4G and G3BP1 were not altered by exogenous AGS3 expression. Oxidative cell stress resulted in the generation of distinct AGS3 BMCs (60 ± 5% of Hela and COS-7 cells with AGS3 BMCs) and SG BMCs (97 ± 3% Hela and COS-7 cells with SG BMCs) that did not exhibit any colocalization as determined by fluorescent microscopy (Figure 2A-B). Similar results were obtained with the endogenous (eIF4G) and transfected (G3BP1) protein markers of SG-BMCs. The AGS3-BMCs observed with cell stress were also distinct from RNA processing BMCs defined by the P-body protein Dcp1a (Supp. Figure 1A).

**Figure 2.**
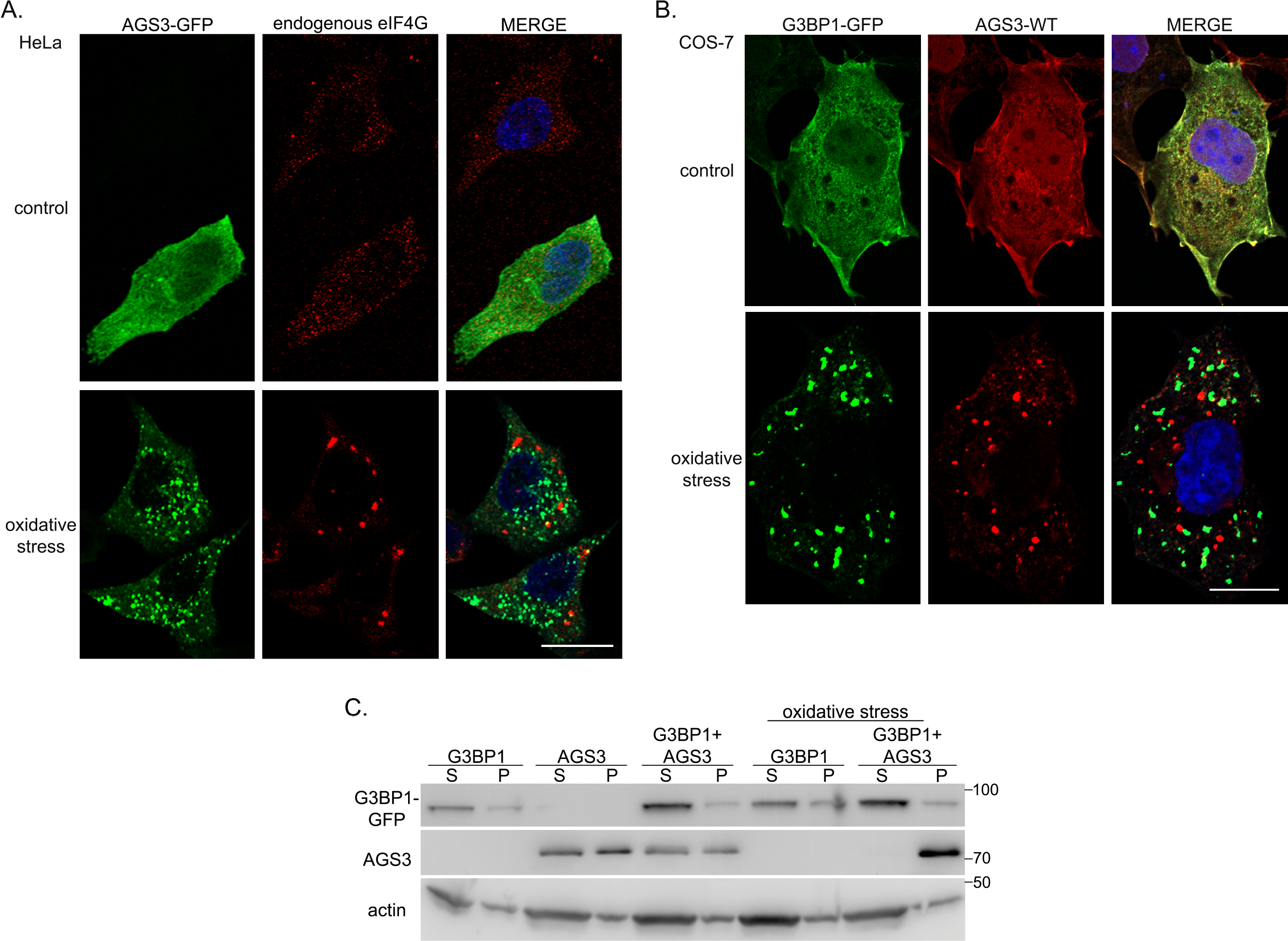
Determination of the distribution of AGS3 and stress granule proteins with and without oxidative cell stress. A) HeLa cells stably expressing AGS3-GFP were exposed to oxidative cell stress (0.5 mM sodium arsenate for 30 min) and processed for immunocytochemistry using eIF4G antibody (1:100 dilution). B) COS-7 cells were transfected with G3BP1-GFP (800 ng) with and without pcDNA3::AGS3 (200 ng). 24 hours following plasmid transfection, cells were exposed to oxidative stress (0.5 mM sodium arsenate for 30 min) and processed for immunocytochemistry using AGS3 (G-2) antibody (1:100 dilution). Fluorescent imaging was performed by confocal microscopy as described in *“Materials and Methods”.* The images are representative of at least five independent experiments and are presented at 63x magnification. C) Cell lysates were fractionated as supernatant (S) and pellet (P) as described in “Materials and Methods” and aliquots processed for sodium dodecyl sulfate – polyacrylamide gel electrophoresis (SDS-PAGE) on precast gels (4-20% gradient gel) and subsequent immunoblotting. Scale bars: 10 μm.

The properties of the AGS3 BMCs in comparison to the SG BMCs were further evaluated by immunoblots of gel transfers following differential fractionation of cell lysates as described in “Experimental Procedures”. The SG BMC protein marker G3BP1 is primarily distributed to the soluble fraction of the cell lysate, and this is not altered by oxidative cell stress (Figure 2C). In contrast, in unstressed cells AGS3 is found in both the cell lysate pellet and soluble fractions (Figures 2C). The distribution of AGS3 to the cell fractions is not altered by cotransfection of G3BP1 consistent with the lack of colocalization of the two proteins by fluorescent microscopy (Figure 2A-B). Furthermore, while oxidative stress does not cause any changes on G3BP1 fractionation, it significantly shifted AGS3 to pellet fraction. These data are consistent with the hypothesis that the stress induced AGS3 BMCs exhibit properties distinct from the SG-BMCs defined with the marker protein G3BP1. A similar fractionation pattern was obtained when AGS3 was coexpressed with the P-body protein marker Dcp1a (Supp. Figure 1B).

### Regulation of stress-induced AGS3 BMCs

As an initial approach to further characterize the AGS3 BMCs generated by oxidative cell stress, we asked two questions. First, is the formation of the stress-induced AGS3 BMCs regulated by protein binding partners involved with signal processing? Second, do the various AGS3 BMCs exhibit different protein diffusion kinetics? As indicated in Figure 3A, oxidative cell stress failed to generate the AGS3 punctate structures when the AGS3 binding partner Gαi3 was co-expressed in the cells with AGS3 (Figure 3A). We also addressed an additional cell stressor involving altered intracellular pH that occurs upon incubation of cells with HBSS in the absence of bicarbonate. Altered pHi also led to the generation of AGS3 BMCs and this was also prevented by co-expression of the AGS3 binding partner Gαi3 (Figure 3A). Initial studies indicate that within the AGS3 BMCs generated in response to cell stress, AGS3 may assemble into a higher order structure through disulfide bridging with other proteins (Supp. Figure 2) based on the differences of migration of the protein through non-reducing, denaturing polyacrylamide gels as compared to the migration pattern observed with standard, reducing denaturing polyacrylamide gel electrophoresis. As observed with the fluorescent microscopic images in Figure 3A, the slower migrating, higher order assembly of AGS3 observed by gel electrophoresis under non-reducing conditions, was also not observed with co-expression of the AGS3 binding partner Gαi3.

**Figure 3.**
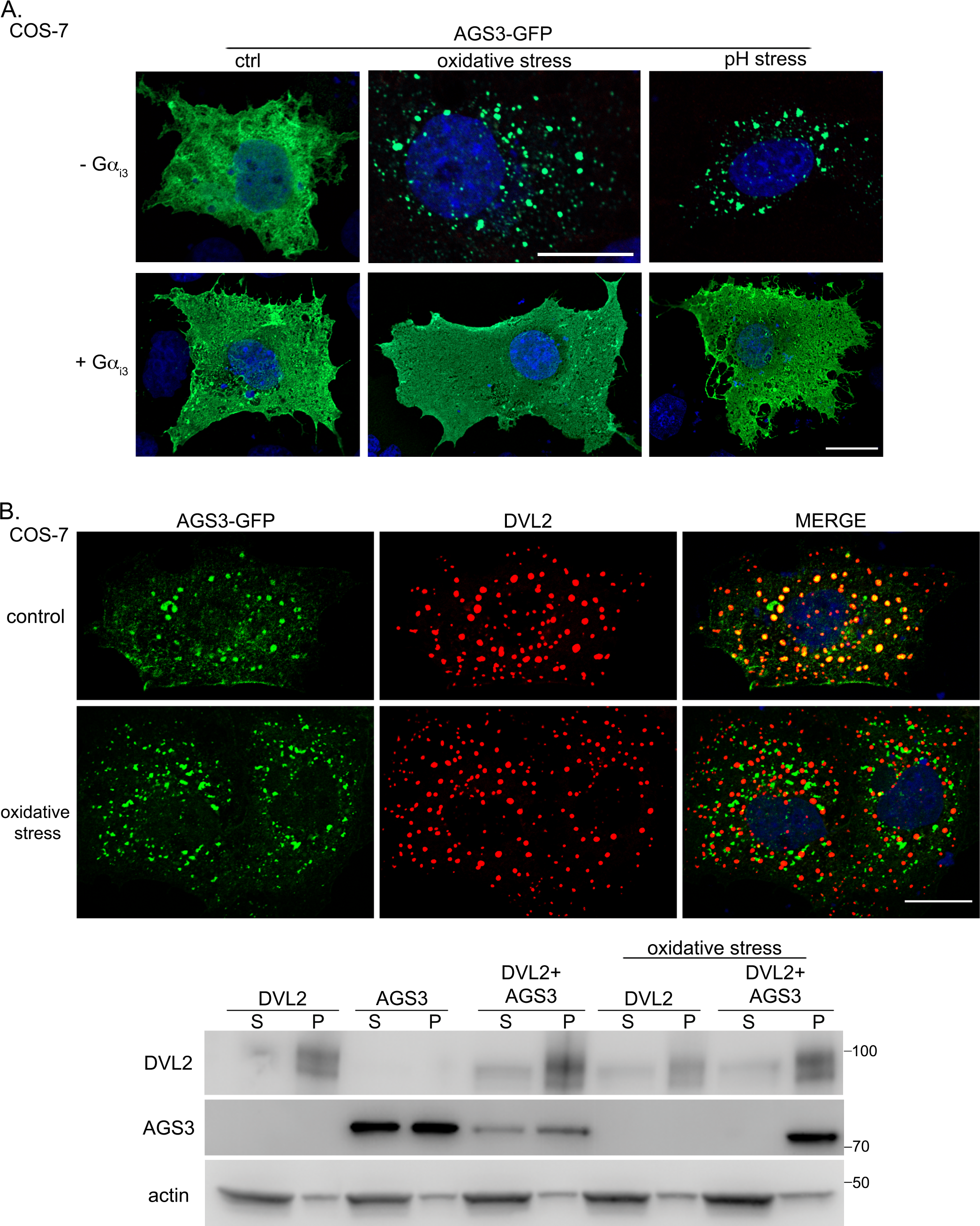
Influence of signal processing proteins on stress-induced AGS3 BMCs. **A) Influence of G-proteins.** COS-7 cells were transfected with pEGFP::AGS3 (250 ng) with and without pcDNA3::Gαi3 (750 ng). 24 hours following plasmid transfection, cells were exposed to oxidative cell stress (0.5 mM sodium arsenate for 30 min) or incubated in Hank’s Balanced Salt Solution (HBSS) without bicarbonate (30 min) to alter intracellular pH and processed for fluorescent microscopy or gel electrophoresis. Fluorescent imaging was performed with confocal microscopy as described in *“Materials and Methods”.* The images are representative of three independent experiments and are presented at 63x magnification. Scale bars: 10 μm. **(B) AGS3-DVL2 biomolecular condensates and oxidative stress.** COS-7 cells were co-transfected with pcDNA3::AGS3 (200 ng) and pRc/CMV2::DVL2 (800 ng) for 24 hours and exposed to oxidative cell stress (0.5 mM sodium arsenate for 30 min) and processed for immunocytochemistry using AGS3 (G-2) (1:100 dilution) and DVL2 (1:100 dilution) antibodies. Fluorescent imaging was performed with confocal microscopy as described in “Materials and Methods”. Images are representative of at least five independent experiments and are presented at 63x magnification. Scale bar: 10 μm. Cell lysates were fractionated as supernatant (S) and pellet (P) as described in “Materials and Methods” and processed for SDS-PAGE (4-20% gradient gel) and immunoblotting.

We also addressed the influence of cell stress on the signaling complex DVL2-AGS3 BMCs (Vural et al., 2018, Vural and Lanier, 2020). In COS7 cells expressing both DVL2 and AGS3, the two proteins appear to colocalize in defined BMCs (Figure 3B) as previously reported (Vural et al., 2018, Vural and Lanier, 2020). This apparent colocalization was disrupted with oxidative cell stress (Figure 3B). Immunoblots of gel transfers subsequent to cell lysis indicated that DVL2 was enriched in the cell lysate pellet and the relative distribution of DVL2 between the supernatant and pellet was not altered by oxidative cell stress (Figure 3B immunoblot). In contrast, oxidative cell stress again shifted AGS3 to the insoluble pellet fraction (Figure 3B) as observed in stress granule and P-body marker coexpression experiments (Figure 2B, Supp. Figure 1B). These data indicate that the stress-induced generation of AGS3 BMCs is regulated by the signaling protein Gαi3, but not by the AGS3 binding partner DVL2.

We next explored the fluidity or rigidity of the stress-induced AGS3 BMCs with respect to the mobility of the AGS3 protein within the cell by evaluating fluorescent recovery of AGS3 signals in different AGS3 BMCs following photobleaching or FRAP. With FRAP, it is inferred that a quick recovery of the fluorescent signal following photobleaching for any specific protein indicates that the protein of interest is “mobile” in the context of the specific microenvironment evaluated. On the other hand, the lack of such recovery of the fluorescent signal indicates a lack of protein mobility suggesting more of a structurally rigid entity or a segregated microenvironment within the cell.

We addressed this question of AGS3 “mobility” by evaluating FRAP for three types of AGS3 BMCs: DVL2-AGS3 BMCs, AGS3-T602A BMCs, AGS3 BMCs induced by oxidative cell stress. Mutation of candidate S/T phosphorylation sites in the GPR domain results in the generation of AGS3 punctate structures and the T602 residue (T625 – G-protein-signaling modulator 1 isoform a, NP_653346) was identified as a key site for this regulation in that mutation of this single residue directs AGS3 to punctate structures in transfected cell systems (Vural et al., 2018). For DVL2-AGS3 BMCs, the recovery of the AGS3 fluorescent signal following photobleaching occurred within 18 seconds indicating a dynamic and fluid BMC with respect to the specific protein AGS3 (Figure 4A - upper panel). In contrast, for the AGS3-T602A BMCs and AGS3 BMCs induced by oxidative cell stress (Figure 4A - middle and lower panels), the recovery of the AGS3 fluorescent signal following photobleaching exhibited markedly different diffusion kinetics (Figure 4B) indicating restricted mobility of AGS3 within these types of BMCs, which is consistent with more of a biophysically rigid BMC as compared to DVL2-AGS3 BMCs.

**Figure 4.**
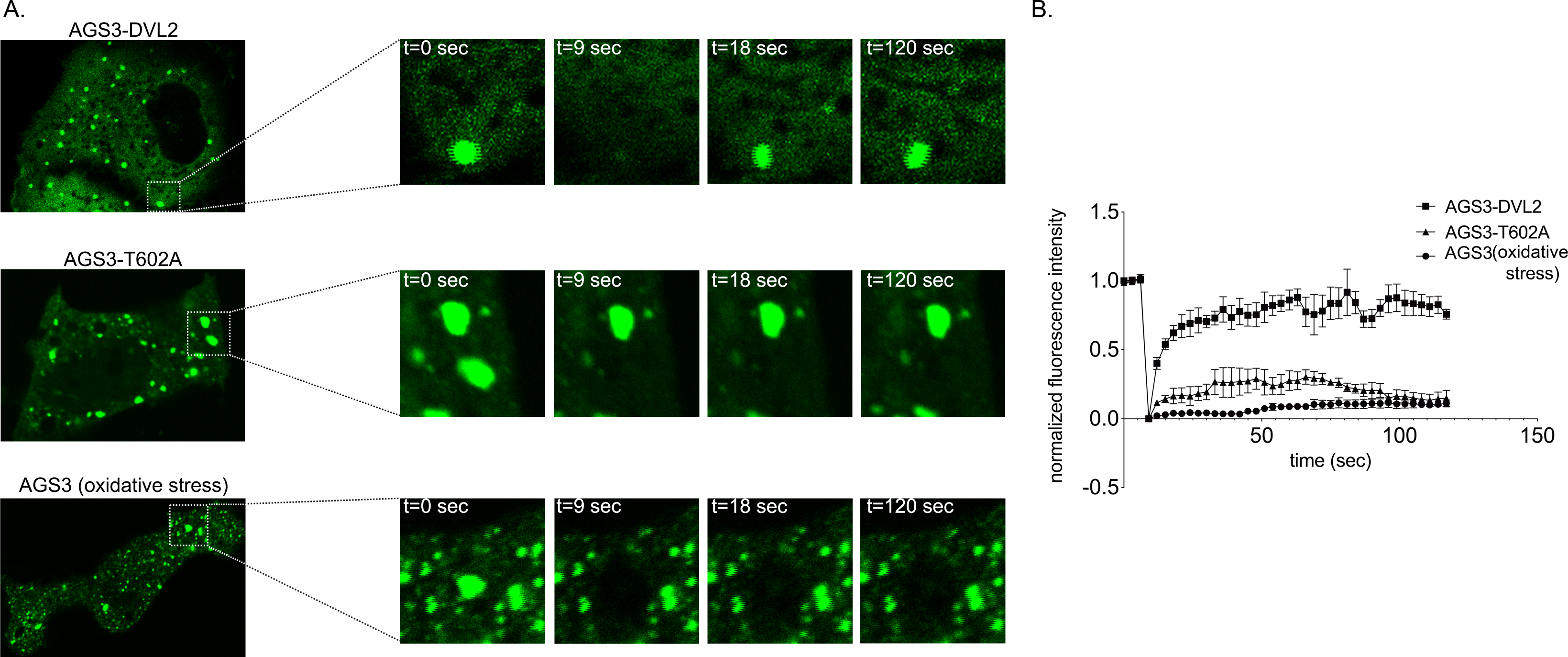
Mobility of AGS3 in the context of defined BMCs. The fluorescent recovery after photobleaching was determined in COS-7 cells as described in “Materials and Methods” for a) AGS3- DVL2 BMCs generated by co-expression of AGS3-GFP (200 ng) and pRc/CMV2::DVL2 (800 ng), b) AGS3-BMCs generated exposed to oxidative cell stress (0.5 mM sodium arsenate for 30 min) and c) AGS3-BMCs observed with the phosphorylation-deficient mutant AGS3-T602A-GFP. Individual AGS3- DVL2, AGS3-T602A and AGS3-GFP puncta induced by oxidative cell stress (n= 5-10) were bleached (t=9 sec) as shown in the represented enlarged insets for each experimental group and fluorescence intensity was measured at every 3 seconds for 2 minutes as described in “Materials and Methods”. B) The fluorescence intensity was normalized and the data presented as the mean intensity ± SEM.

## DISCUSSION

While AGS3 is implicated in a number of diverse cell and system functions and the biochemistry of AGS3 with respect to structural domains and its interaction with G-proteins is well-investigated, the mechanisms by which the biochemistry of AGS3 signaling is processed and regulated with respect to cell and tissue function is not clear. Similar challenges exist for various signaling transduction systems in that there is simply a level of complexity in terms of “translating” the biochemistry of signaling systems (e.g. protein interactions, signal transduction networks) into specific and defined human phenotypes. This has presented challenges to both understanding disease mechanisms and the development of new therapeutic modalities for any given disease. Toward this end and with the enabling technologies advanced over the last several years with respect to bio-imaging and protein dynamics within cells, we explored the possibility of using image phenotype profile screens to gain further insight into signal-function processes for the family of AGS proteins (Vural et al., 2010, Oner et al., 2010, Oner et al., 2013b, Vural et al., 2016, Vural et al., 2018, Vural and Lanier, 2020). This concept of “image-signaling system connectivity” was recently illustrated with the discovery of the AGS3 interaction with the disheveled signaling nexus (Vural and Lanier, 2020). In the current manuscript, we further expanded this platform to examine the positioning of AGS3 with respect to other types of functional punctate structures in the cell and if this positioning was regulated by cell stress and signal transducing proteins.

Given the cell biology and protein dynamics of AGS3, its propensity to exist in defined and distinct punctate structures within the cell, and the regulated generation of such punctate structures, we have refer to these punctate structures as AGS-BMCs,. BMCs are non-membranous and organelle-sized (i.e., micron-scale) structures formed by liquid-liquid phase separations (Banani et al., 2017). By spatially organizing and coordinating the physical assembly of a specific subset of molecules sequestered from the rest of the cytoplasm, BMCs may have a range of affects on the activity/function of individual proteins or assemblies of various related proteins including a) enhanced activity through concentrating enzymes and substrates; b) reduced activity through sequestration; and c) modulated activity through exclusion of regulatory factors (Huang et al., 2016, Case et al., 2019, Lyon et al., 2021, Peeples and Rosen, 2021). Additionally, BMCs exhibit extraordinary sensitivity to solution conditions such as oxidative stress, temperature, pH, and ionic strength, where biological systems exploit them as a mechanism to sense potentially deleterious environmental conditions and induce appropriate homeostatic responses (Franzmann and Alberti, 2019). These core properties of BMCs may be operational within a wide range of physiological (transcription and genome organization, immune response, neuronal synaptic signaling) and pathological events (neurodegeneration, cancer and viral infections) within the cell (Wang et al., 2021).

Thus, there are two central concepts advanced with this manuscript and our recent work in this area. The first concept is that AGS3 can engage with defined BMCs in a regulated manner. The second concept is that AGS3, given its regulated interaction with key cell signaling proteins, is a core element of a new type of BMCs that serve as previously unappreciated signal processing nodes. Indeed, AGS3 exhibits an inherent propensity to form distinct, non-membranous punctate structures, which do not appear to associate with any defined vesicle or organelle markers or be enveloped by a lipid bilayer (Vural et al., 2010, Vural et al., 2018). The linker and GPR regions of AGS3 protein exhibit high intrinsic disorder and a similarly strong intrinsic disorder pattern is observed with the Group II AGS proteins AGS4/GPSM3 and AGS5/GPSM2) (Uversky, 2014). Of note, BMCs are suggested to be involved in the role of AGS5/GPSM2 in actin-bundling in stereocilia that may directly relate to the role of this protein in Chudley-McCullough syndrome (Shi et al., 2022).

As reported herein, the generation of AGS3 BMCs and SG BMCs was markedly increased by oxidative cell stress. However, and of particular note, the AGS3 and SG BMCs generated in response to oxidative cell stress were distinct from each other and exhibited different biochemical properties. Upon cell lysis, in non-stressed cells, AGS3 distributes to both a cytosolic supernatant and a membrane pellet fraction, but with oxidative stress AGS3 is primarily found in the membrane pellet. In contrast, oxidative stress did not alter the relative distribution of DVL2, the SG BMC protein G3BP1, or the P-body protein Dcp1a in the pellet and supernatant subsequent to cell lysis. The AGS3-BMC induced by stress is distinct from the population of stress granules as defined by the protein marker G3BP1. However, there are likely distinct subpopulations of stress granules that differ in their composition, biophysical properties, and their functional role within the stress granule ecosystem of the cell response to stressors (Jain et al., 2016, Shiina, 2019, Cirillo et al., 2020, Marmor-Kollet et al., 2020). Of note AGS3 (GPSM1) appears enriched in a subpopulation of stress granules defined with the marker proteins FMR1/FXR1 (Marmor-Kollet et al., 2020).

These data are generally consistent with the concept that AGS3 is capable of defining a distinct population of BMCs. One working hypothesis is that the dynamics of structural organization of AGS3 allows a degree of microfluidity for the protein within in the cell. Two specific observations from the current study have particular relevance to this hypothesis: 1) the altered fluidity of AGS3 BMCs as compared to AGS3-DVL2 BMCs as determined by FRAP and 2) the altered migration of immunoreactive AGS3 species as visualized with immunoblots following non-reducing, denaturing polyacrylamide gel electrophoresis. Both oxidative cell stress and altered cellular pH resulted in the visualization of immunoreactive AGS3 species of higher apparent Mr in the membranous pellet fraction isolated following cell lysis. Such a migration pattern may represent dimerization or oligomerization of AGS3 protein via intermolecular disulfide bonds or stress-induced rearrangements of structural domains within AGS3 that influence coordination of posttranslational events and protein modifications (Niwa et al., 2007). The prevention of the generation of AGS3 BMCs by Gαi co-expression further suggests that the direct interaction of Gαi with AGS3 stabilizes specific conformations of the protein.

Of particular note, both the AGS3-BMCs generated by oxidative cell stress and the punctate structures observed with a phospho-deficient AGS3 (AGS3-T602A) do not appear to colocalize with DVL2 upon co-expression as is the case for AGS3 in non-stressed cells (Vural et al., 2018). These data suggest that the DVL2-AGS3 BMCs are distinct from the BMCs observed with phospho-deficient AGS3 (AGS3- T602A) and with oxidative cell stress. The FRAP studies reported in this study indicated that this was indeed the case as the AGS3 within the AGS3-DVL2 BMCs clearly existed in more of a fluid state as compared to the AGS3 that localized in the BMCs generated by oxidative cell stress and those observed with the phospho-deficient AGS3 (AGS3-T602A). These data indicate an ongoing internal rearrangement of proteins within AGS3-DVL2 puncta as well as an in-and-out trafficking of AGS3-GFP molecules to and from the puncta suggesting they exist in a dynamic and reversible state. AGS3-DVL2 puncta possess a higher mobility and fusion rate with fluid-like characteristics. On the other hand, the lack or a very-low degree of fluorescence recovery of AGS3 BMCs generated by oxidative cell stress and with phospho-deficient AGS3-T602A suggests a gel-like and less fluid texture. Indeed, the AGS3-BMCs generated by oxidative cell stress were still observed up to 24 hours after removal of the cell stressor (Vural, A and Lanier, SM - unpublished observations). Based upon this series of results and current concepts in the BMC field, the fluid association of AGS3 with DVL2 BMCs and the reduced fluidity of AGS3 in the stress-induced AGS3 BMCs may reflect, respectively, “client” and “scaffolding” roles for AGS3 (Banani et al., 2016, Espinosa et al., 2020, Lyon et al., 2021).

The results of these studies indicate that AGS3 is associated with different signal processing BMCs and that this association is regulated by cell stress and the signaling proteins Gαi3 and DVL2. A further understanding of the life course and molecular composition of AGS3 BMCs may provide unexpected insight into the range of pathophysiologies and system functions associated with AGS3 and related Group II AGS proteins.

## Acknowledgements

AV gratefully acknowledges the support of Dr. Raymond Mattingly (Chair, Department of Pharmacology, School of Medicine, Wayne State University, Detroit, MI) for his support and encouragement. SML also acknowledges and is greatly appreciative of the opportunity and support provided by M. Roy Wilson (President, Wayne State University) during his tenure at Wayne State University in Detroit, MI. Cell imaging studies were conducted through the Microscopy, Imaging & Cytometry Resources Core at Wayne State University, Detroit, MI. As always, SML appreciates the valued suggestions and gracious engagement of the many students, fellows, colleagues, and collaborators that have contributed to the body of work involving Activators of G protein signaling since their initial discovery.

---

Supplemental Figure 1. D**e**termination **of the distribution of AGS3 and P-body proteins with and without oxidative cell stress.** A) HeLa cells stably expressing AGS3-GFP were exposed to oxidative cell stress (0.5 mM arsenate for 30 min) and processed for immunocytochemistry using Dcp1a antibody (1:100 dilution). The images are representative of three independent experiments and are presented at 63x magnification. B) COS-7 cells were transfected with Dcp1a-GFP (800 ng) with and without pcDNA3::AGS3 (200 ng). After 24 hours, cells were exposed to oxidative stress (0.5 mM sodium arsenate for 30 min) and processed for immunoblotting by fractionating the cell lysates as supernatant (S) and pellet (P) and SDS-PAGE (4-20% gradient gel) as described in “Materials and Methods”. Scale bar: 10 μm

Supplemental Figure 2. I**n**fluence **of G-proteins on biochemical properties of AGS3-BMCs.** COS-7 cells were transfected with pEGFP::AGS3 (250 ng) with and without pcDNA3::Gαi3 (750 ng). 24 hours following plasmid transfection, cells were exposed to oxidative cell stress (0.5 mM sodium arsenate for 30 min) or incubated in Hank’s Balanced Salt Solution (HBSS) without bicarbonate (30 min). Following cellular lysis, samples were processed for immunoblotting described in “Materials and Methods” without and with added reducing agent and with protein separation by non-reducing SDS-PAGE (4-20% gradient gel). S – supernatant, P – pellet.

