## Supplementary figures and images for "Biomolecular Condensates defined by Receptor Independent Activator of G protein Signaling: Properties and Regulation"

### Supplemental Figures 1&2

Supplemental Figure 1.

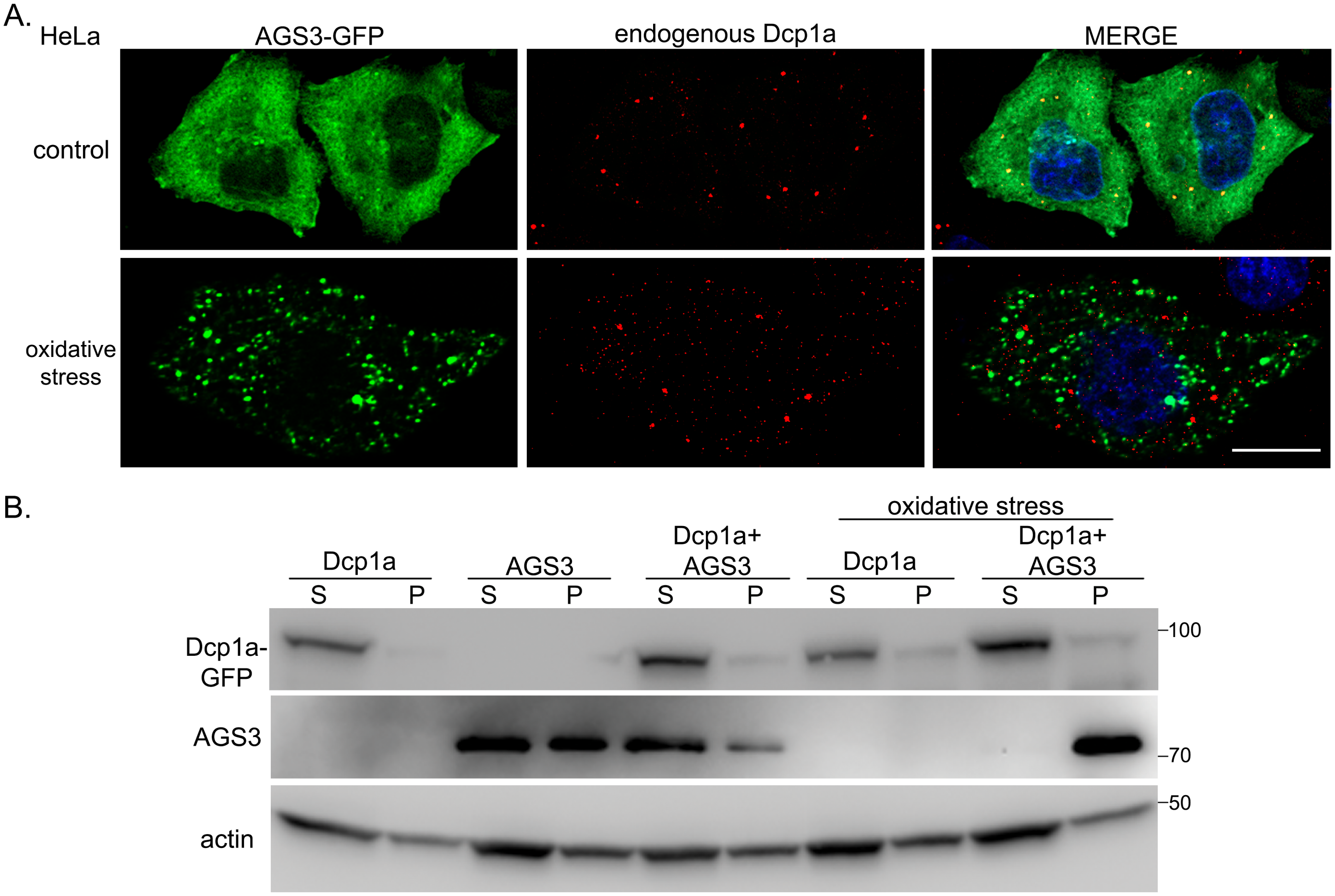

Supplemental Figure 2.

A.

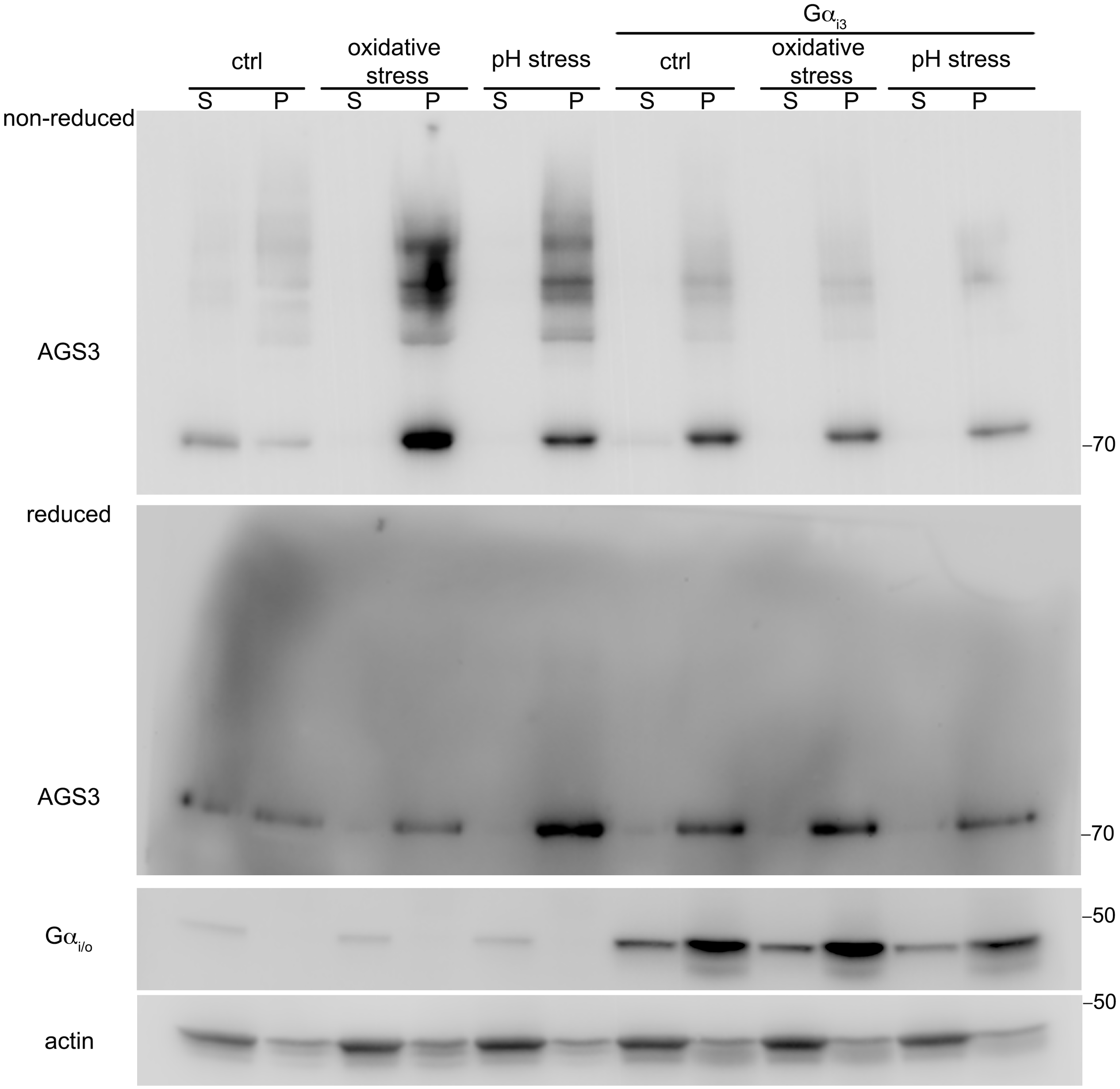
